# Partial agonism improves the anti-hyperglycaemic efficacy of an oxyntomodulin-derived GLP-1R/GCGR co-agonist

**DOI:** 10.1101/2021.03.02.433157

**Authors:** Phil Pickford, Maria Lucey, Roxana-Maria Rujan, Emma Rose McGlone, Stavroula Bitsi, Fiona Ashford, Ivan R Corrêa, David J Hodson, Alejandra Tomas, Giuseppe Deganutti, Christopher A Reynolds, Bryn M Owen, Tricia M Tan, James Minnion, Ben Jones, Stephen R Bloom

**Author notes:** Corresponding author: Ben Jones.

## Abstract

**Objective:** Glucagon-like peptide-1 and glucagon receptor (GLP-1R/GCGR) co-agonism can maximise weight loss and improve glycaemic control in type 2 diabetes and obesity. In this study we investigated the cellular and metabolic effects of modulating the balance between G protein activation and β-arrestin-2 recruitment at GLP-1R and GCGR using oxyntomodulin (OXM)-derived co-agonists. This strategy has been previously shown to improve the duration of action of GLP-1R mono-agonists by reducing target desensitisation and downregulation.

**Methods:** Dipeptidyl dipeptidase-4 (DPP-4)-resistant OXM analogues were generated and assessed for a variety of cellular readouts. Molecular dynamic simulations were used to gain insights into the molecular interactions involved. In vivo studies were performed in mice to identify effects on glucose homeostasis and weight loss.

**Results:** Ligand-specific reductions in β-arrestin-2 recruitment led to reduced GLP-1R internalisation and prolonged glucose-lowering action *in vivo*. The putative benefits of GCGR agonism were retained, with equivalent weight loss compared to the GLP-1R mono-agonist liraglutide in spite of a lesser degree of food intake suppression. The compounds tested showed only a minor degree of biased agonism between G protein and β-arrestin-2 recruitment at both receptors and were best classified as partial agonists for the two pathways measured.

**Conclusions:** Diminishing β-arrestin-2 recruitment may be an effective way to increase the therapeutic efficacy of GLP-1R/GCGR co-agonists. These benefits can be achieved by partial rather than biased agonism.

## 1 Introduction

Insulin and glucagon are traditionally viewed as opposing protagonists in the hormonal control of blood glucose. Pharmacological approaches to potentiate glucose-stimulated insulin secretion (GSIS), such as analogues of the incretin glucagon-like pepide-1 (GLP-1), have been successfully exploited over many years for the treatment of type 2 diabetes (T2D) [1]. However, decades of attempts to develop glucagon receptor (GCGR) antagonists for clinical use have so far failed to yield any approved therapeutic agents [2]. A significant problem appears to be the development of hepatic steatosis [3–6]. Contrasting with this traditional approach, GCGR agonism has emerged as a credible component of combined therapeutic strategy for the treatment of obesity and T2D in which GLP-1R and GCGR are concurrently targeted [7, 8], thereby recapitulating the effects of the endogenous GLP-1R/GCGR co-agonist oxyntomodulin (OXM) [9]. Well established effects of glucagon on energy expenditure [10], leading to enhanced weight loss and ultimately improvements in insulin sensitivity [11], might then be safely realised in the context of GLP-1R-mediated protection against acute hyperglycaemia. Glucagon is also insulinotropic, an effect which derives from action at both GLP-1R and GCGR [12, 13].

Biased agonism is a concept in which different ligands for the same receptor selectively couple to different intracellular effectors [14], potentially providing a means to improve their therapeutic window by reducing activation of pathways associated with adverse effects [15]. For G protein-coupled receptors (GPCRs), bias is commonly, but not always, expressed as a relative preference for recruitment of G proteins *versus* β-arrestins, i.e. two of the most proximal interactors recruited to the activated receptor, as well as their corresponding signalling intermediates. Both GLP-1R and GCGR are primarily coupled to cAMP generation through Gα_s_ activation, with recruitment of β-arrestins being associated with receptor desensitisation, endocytosis and diminished long term functional responses [16, 17]. Whilst the therapeutic benefits of biased GLP-1R agonism have been demonstrated in preclinical studies on a number of occasions [18–21], the application of this principle to GLP-1R/GCGR co-agonism is less explored. A recent study reported bias profiles for a selection of investigational dual GLP-1R/GCGR agonists, but it is not clear what role bias plays in their metabolic effects [22].

In this study we aimed to develop GLP-1R/GCGR co-agonists with altered signalling properties but otherwise equivalent characteristics, which might then be used to assess the functional impact of bias *in vitro* and *in vivo*. Focussing on the peptide N-terminus, we evaluated dipeptidyl dipeptidase-4 (DPP-4)-resistant peptides featuring 2-aminoisobutyric acid (AIB) at position 2, in combination with which a switch between glutamine (Q) and histidine (H) at position 3 was able to alter the maximum responses (i.e. efficacy) for G protein and β-arrestin recruitment, to varying degrees, at both receptors. Molecular dynamics simulation of glucagon analogues interacting with GCGR was used to gain insight into the molecular interactions underlying these differences. By comparing the metabolic effects of a pair of matched peptides with these sequence substitutions, we demonstrate that reduced recruitment efficacy of β-arrestins translates to improved efficacy in preclinical rodent models of obesity, consistent with a similar effect previously observed for GLP-1R mono-agonists [18–21]. Our study therefore suggests a viable strategy to optimise GLP-1R/GCGR co-agonism for enhanced therapeutic efficacy.

## 2 Materials and methods

### 2.1 Peptides

All peptides were obtained from Wuxi Apptec and were of at least 90% purity.

### 2.2 Cell culture

HEK293T cells were maintained in DMEM supplemented with 10% FBS and 1% penicillin/streptomycin. PathHunter CHO-K1-βarr2-EA cells stably expressing human GLP-1R, GCGR or GIPR, and PathHunter CHO-K1-βarr1-EA cells stably expressing GCGR, were obtained from DiscoverX and were maintained in Ham’s F12 medium with 10% FBS and 1% penicillin/streptomycin. Stable HEK293-SNAP-GLP-1R cells [23] were maintained in DMEM supplemented with 10% FBS, 1% penicillin/streptomycin and 1 mg/ml G418.

### 2.3 Animal husbandry

Animals were maintained in specific pathogen-free facilities, with *ad lib* access to food (except prior to fasting studies) and water. Studies were regulated by the UK Animals (Scientific Procedures) Act 1986 of the U.K. and approved by the Animal Welfare and Ethical Review Body of Imperial College London (Personal Project License PB7CFFE7A) or University of Birmingham (Personal Project License P2ABC3A83). Specific procedures are described below.

### 2.4 Primary islet isolation and culture

Mice were euthanised by cervical dislocation before injection of collagenase solution (1 mg/ml, Serva NB8, or S1745602, Nordmark Biochemicals) into the bile duct. Dissected pancreata were then digested for 12 min at 37°C in a water bath before purification of islets using a Ficoll (1.078) or Histopaque (Histopaque-1119 and −1083) gradient. Islets were hand-picked and cultured (5% CO_2_, 37°C) in RPMI medium containing 10% FBS and 1% penicillin/streptomycin.

### 2.5 Primary hepatocyte isolation and culture

Hepatocytes from adult male C57Bl/6J mice were isolated using collagenase perfusion [24]. After filtering and washing, cells were used directly for assaying of cAMP responses as described below.

### 2.6 NanoBiT assays and calculation of bias between mini-G_s_ and β

The assay was performed as previously described [21]. HEK293T cells in 12-well plates were transfected with 0.5 µg GLP-1R-SmBiT plus 0.5 µg LgBiT-mini-G_s_ [25] (a gift from Prof Nevin Lambert, Medical College of Georgia), or with 0.05 μg GLP-1R-SmBiT, 0.05 μg LgBiT-β-arrestin-2 (Promega) plus 0.9 µg pcDNA3.1 for 24 hours. Cells were detached with EDTA, resuspended in HBSS, and furimazine (Promega) was added at a 1:50 dilution from the manufacturer’s pre-prepared stock. After dispensing into 96-well white plates, a baseline read of luminescent signal was serially recorded over 5 min using a Flexstation 3 instrument at 37°C before addition of the indicated concentration of ligand, after which signal was repeatedly recorded for 30 min. For AUC analysis, results were expressed relative to individual well baseline for AUC calculation over the 30-min stimulation period. Baseline drift over time frequently led to a negative AUC for vehicle treatment, which was subtracted from all results before construction of 3-parameter curve fits of concentration-response in Prism 8.0. Bias was calculated using two approaches. Firstly, the log max/EC_50_ method [26] was used, with the ratio of E_max_ to EC_50_ from 3-parameter fits for each ligand used to quantify agonism. After log_10_ transformation, responses were expressed relative to the reference agonist on a per assay basis to give Δlog(E_max_/EC_50_) for each pathway. Pathway-specific values were then expressed relative to each other to give ΔΔlog(E_max_/EC_50_), i.e. the log bias factor. Secondly, a method derived from kinetic curve fitting was used [27]. Here, kinetic responses for a single maximal agonist concentration were normalised at each time-point to the vehicle response prior to curve fitting. Mini-G_s_ responses were fitted using the one-phase exponential association equation in Prism 8.0. β-arrestin-2 responses were fitted using the biexponential equation described in [27]. The agonist efficacy term *k*_*τ*_ was derived from these data as described [27] for each agonist and, after log_10_ transform, the SRB103Q response was expressed relative to SRB103H as the reference agonist on a per assay basis to give Δlog *k*_*τ*_. Pathway-specific values were then expressed relative to each other to give ΔΔlog *k*_*τ*_, i.e. the log bias factor.

### 2.7 Biochemical measurement of cAMP production

PathHunter cells were stimulated with the indicated concentration of agonist for 30 min at 37°C in serum free medium, without phosphodiesterase inhibitors. Primary dispersed mouse islet cells, prepared by triturating intact islets in 0.05% trypsin/EDTA for 3 min at 37°C, were stimulated with indicated concentration of agonist for 5 min at 37°C in serum free medium, with 11 mM glucose and 500 µM IBMX. Primary mouse hepatocytes were stimulated in serum free medium with 100 µM IBMX for 10 min, or for 16 hours in serum free medium with 100 µM IBMX added for the final 10 min of the incubation. At the end of each incubation, cAMP was then assayed by HTRF (Cisbio cAMP Dynamic 2) and concentration-response curves constructed with 3-parameter curve fitting in Prism 8.0.

### 2.8 Dynamic cAMP imaging in intact islets

C57Bl/6 (n = 7) or Ins1^tm1.1(cre)Thor+/-^ (n = 2) mice were used as islet donors and were phenotypically indistinguishable. Islets were transduced with epac2-camps for 48 hours using an adenoviral vector (a kind gift from Prof Dermot M. Cooper, University of Cambridge). Epac2-camps is well validated, relatively pH insensitive, and senses cAMP concentrations in the ranges described in islets [28, 29]. Dynamic cAMP imaging was performed as previously described [30] using a Crest X-Light spinning disk system, coupled to a Nikon Ti-E microscope base and 10× objective. Excitation was delivered at λ = 430-450 nm using a SPECTRA X light engine. Emitted signals were detected using a 16-bit Photometrics Evolve Delta EM-CCD at λ = 460-500 nm and 520-550 nm for cerulean and citrine, respectively. For imaging, islets were maintained in HEPES-bicarbonate buffer (pH 7.4) containing (in mM): 120 NaCl, 4.8 KCl, 24 NaHCO_3_, 0.5 Na_2_HPO_4_, 5 HEPES, 2.5 CaCl_2_, 1.2 MgCl_2_, 16.7 *D*-glucose. The experiment was conducted to determine responses to agonist-naïve islets (“acute”), or with a “rechallenge” design, in which islets were first treated for 4 hours with 100 nM agonist followed by washout (2 washes, 30 min) before commencing imaging. During the imaging, islets were stimulated with 100 nM agonist for 15 min, starting at T=5 min, followed by application of 10 µM forskolin as a positive control. FRET responses were calculated as the fluorescence ratio of Cerulean/Citrine and normalized as F/F_0-5_, where F denotes fluorescence at any given time point and F_0-5_ denotes average fluorescence during 0-5 mins.

### 2.9 High content imaging assay for receptor internalisation

The assay was adapted from a previously described method [31]. HEK293T cells were seeded in black, clear bottom, plates coated with 0.1% poly-D-lysine, and assayed 24 hours after transfection with SNAP-tagged GLP-1R or GCGR plasmid DNA (0.1 µg per well). Cells were labelled with the cleavable SNAP-tag probe BG-S-S-549 (a gift from New England Biolabs) in complete medium for 30 min at room temperature. After washing, fresh, serum-free medium ± agonist was added. At the end of the incubation, medium was removed, and wells were treated with for 5 min at 4°C with Mesna (100 mM, in alkaline TNE buffer, pH 8.6) to remove BG-S-S-549 bound to residual surface receptor without affecting the internalised receptor population, or with alkaline TNE buffer alone. After washing, cellular imaging and processing by high content analysis was performed as previously described [31] to quantify the amount of internalised receptor from fluorescence intensity readings.

### 2.10 Preparation and imaging of fixed cell samples to observe receptor internalisation

Cells were seeded onto coverslips coated with 0.1% poly-D-lysine and assayed 24 hours after transfection with SNAP-tagged GLP-1R or GCGR plasmid DNA (0.5 µg per well of a 24-well plate). Surface labelling of SNAP-tagged GLP-1R was performed using 0.5 µM of the indicated SNAP-Surface probe for 30 min at 37°C before washing with HBSS. Ligands were applied in Ham’s F12 media at 37°C. For fixation, 4% paraformaldehyde (PFA) was applied directly to the medium for 15 min before washing with PBS. Slides were mounted in Prolong Diamond antifade with DAPI and allowed to set overnight. Widefield epifluorescence imaging was performed using a Nikon Ti2E custom microscope platform via a 100× 1.45 NA oil immersion objective, followed by Richardson-Lucy deconvolution using DeconvolutionLab2 [32].

### 2.11 Measurement of GLP-1R internalisation by DERET

The assay was performed as previously described [21]. HEK-SNAP-GLP-1R cells were labelled using 40 nM SNAP-Lumi4-Tb in complete medium for 60 min at room temperature. After washing, cells were resuspended in HBSS containing 24 µM fluorescein and dispensed into 96-well white plates. A baseline read was serially recorded over 5 min using a Flexstation 3 instrument at 37°C in TR-FRET mode using the following settings: λ_ex_ 340 nm, λ_em_ 520 and 620 nm, auto-cutoff, delay 400 µs, integration time 1500 µs. Ligands were then added, after which signal was repeatedly recorded for 30 min. Fluorescence signals were expressed ratiometrically after first subtracting signal from wells containing 24 µM fluorescein but no cells. Internalisation was quantified as AUC relative to individual well baseline, and concentration-response curves generated with Prism 8.0.

### 2.12 DPP-4 peptide degradation assay

10 nmol SRB103Q, SRB103H or GLP-1 were dissolved in 750 µl DPP-4 buffer (100 mM Tris-HCl; pH 8). 10 mU recombinant DPP-4 (R&D Systems), or no enzyme as a control for non-enzymatic degradation over the same time period, was added to the reconstituted peptide. Reactions were incubated at 37°C and 120 µl samples were collected from the reaction vessel at the indicated time-points. 5 µl 10% trifluoroacetic acid (TFA) was added to each sample to terminate enzyme activity. Samples were analysed by reverse-phase high-performance liquid chromatography (HPLC) with a linear acetonitrile/water gradient acidified with 0.1% TFA on Phenomenex Aeris Peptide 3.6 μm XB-C18 Column (150 x 4.6 mm). The eluted peptides were detected at 214 nm. Degradation of peptide was calculated by comparing the area under the peak of the original compound with and without enzyme.

### 2.13 In vivo studies

Lean male C57Bl/6 mice (8-10 weeks of age, body weight 25-30 g, obtained from Charles River) were maintained at 21-23°C and 12-hour light-dark cycles. *Ad libitum* access to water and normal chow (RM1, Special Diet Services), or diet containing 60% fat to induce obesity and glucose intolerance (D12492, Research Diets) for a minimum of 3 months before experiments, was provided. Mice were housed in groups of 4, except for food intake assessments and the chronic administration study, when they were individually caged with 1 week of acclimatisation prior to experiments. Treatments were randomly allocated to groups of mice matched for weight.

### 2.14 Intraperitoneal glucose tolerance tests

Mice were fasted for at least 4 hours before commencing the glucose tolerance test, depending on the peptide treatment length. Mice were injected into the intraperitoneal (IP) cavity with peptide or vehicle (0.9% saline) either 8 hours before, 4 hours before, or at the same as the glucose challenge (acute). Glucose was dosed at 2 g/kg body weight. Blood glucose levels were measured before glucose challenge, then at the times as indicated in the figure using a hand-held glucose meter (GlucoRx® Nexus). Blood samples for insulin were collected at 10 minutes into lithium heparin-coated microvette tubes (Sarstedt, Germany), followed by centrifugation (10,000 RPM, 8 min, 4°C) to separate plasma. Plasma insulin was measured using the Cisbio mouse insulin HTRF kit.

### 2.15 Insulin tolerance tests

Mice were fasted for 2 hours before IP injection of peptide or vehicle (0.9% saline). 4 hours later, baseline blood glucose was taken before recombinant human insulin (Sigma, USA) (0.5 U/kg – 1 U/kg) was injected IP and blood glucose measured 20, 40 and 60 minutes after insulin injection.

### 2.16 Feeding studies

Mice were fasted overnight before the study. Diet was returned to the cage 30 min after IP injection of agonist, with cumulative intake determined by weighing.

### 2.17 Pharmacokinetic study

Mice were administered 0.5 mg/kg peptide via IP injection. 4 hours after injection, blood was acquired by venesection into lithium heparin-coated microvette tubes (Sarstedt, Germany). Plasma was separated by centrifugation at 10,000 g for 8 minutes at 4°C. Plasma concentrations were assessed by radioimmunoassay using an in-house assay as previously described [33], using standard curves generated from each SRB103 peptides to ensure equivalent recovery was obtained.

### 2.18 Chronic administration study

SRB103 peptides were made up in aqueous ZnCl_2_ solution to a molar ratio of 1.2:1 (ZnCl_2_:peptide). Liraglutide (Novo Nordisk) was diluted in sterile water. DIO mice received daily subcutaneous (s.c.) injections of each treatment or vehicle (matched ZnCl_2_) with the dose increased during the first week as indicated on the figure. Body weight and food intake was measured periodically, with food and water available ad libitum. The end-of-study glucose tolerance test was performed 8 hours after the final peptide dose, with mice fasted for 5 hours.

### 2.19 Statistical analysis of biological data

Quantitative data were analysed using Prism 8.0 (GraphPad Software). In cell culture experiments, technical replicates were averaged so that each individual experiment was treated as one biological replicate. Dose responses were analysed using 3 or 4-parameter logistic fits, with constraints imposed as appropriate. Bias analyses were performed as described in Section 2.6. Statistical comparisons were made by t-test or ANOVA as appropriate, with paired or matched designs used depending on the experimental design. Mean ± standard error of mean (SEM), with individual replicates in some cases, are displayed throughout, with the exception of bias analyses for which 95% confidence intervals are shown to allow straightforward identification of biased ligands, for which the 95% confidence bands do not cross zero. Statistical significance was inferred if p<0.05, without ascribing additional levels of significance.

### 2.20 Systems preparation, equilibration and molecular dynamics simulation

We performed molecular dynamics simulations on the active structure of GCGR in complex with peptides GCG, GCG-AIB2 and GCG-AIB2H3, and the C-terminal helix 5 of the α-subunit of the Gs protein. The structure was modelled using MODELLER software (https://salilab.org/modeller) [34]. The templates used were the full-length crystal structure of a partial activated GCGR in complex with NNC1702 peptide (PDB: 5YQZ) [35] and the cryo-EM structure of the active glucagon-like peptide-1 receptor (GLP-1R) (PDB: 6B3J) [36]. Maestro software (https://www.schrodinger.com) was employed to add the missing residue H^1^ and substitute the adequate residues to generate GCG, GCG-AIB2 and GCG-AIB2H3. Once the three systems were complete and the hydrogens added, each system was embedded in a phospholipidic membrane and solvated. The membrane model used was 1-palmitoyl,2-oleoyl-sn-glycero-3-phosphochholine (POPC), which was generated by CHARMM-GUI (http://charmm-gui.org). The simulation box dimensions of the resulting systems was 90 x 90 x 170 A in X, Y and Z directions, respectively. General charge neutrality was obtained by adding neutralizing counter ions Na^+^ and Cl^-^. Each system was subjected to 10 000 cycles of energy minimization in order to eliminate steric clashes and relax the side chains. The final step before running the simulations was represented by the equilibration of the systems, which includes re-orientations of the water and lipid molecules around the protein. The systems were both equilibrated and simulated in an NVT ensemble with semi-isotropic pressure scaling with a constant surface tension dynamic of 0 dyne/cm (through interfaces in the XY plane). The target pressure of 1 bar was achieved using the Monte Carlo barostat, while the target temperature of 300 K was regulated using Langevin dynamics with a collision frequency of 1 ps^-1^. The SHAKE algorithm was used to constrain the lengths of bonds comprising hydrogen atoms. Every system was equilibrated for 32 ns at a time step of 2 fs, then run in 3 replicas for approximately 2 μs, at a time step of 4 fs, using the AMBER force field implemented in the AMBER software package (http://ambermd.org) [37].

### 2.21 MD trajectories analysis

Every replica of a system was merged and aligned on the initial frame using MDTraj (www.mdtraj.org) and then subjected to analysis. The hydrogen bonds and the van der Waals interactions between peptides and receptors were computed using the GetContacts package (https://getcontacts.github.io). The contacts were plotted on the PDB coordinates using in-house scripts and Chimera software (www.cgl.ucsf.edu). The distances between T369^6.60^, located at the top of TM6, and the origin of the cartesian coordinates (0, 0, 0) were quantified by employing the open-source, community-developed library PLUMED 2.0 (www.plumed.org). Using the data provided by PLUMED we further calculated the distribution of the distances by implementing an in-house script. Principal component analysis (PCA) was conducted on the Cα atoms using the R package Bio3D (www.thegrantlab.org) [38]. Prior to PCA we carried out a trajectory frame superposition on Cα atoms of residues 133 to 403 (TM domain) to minimize the root mean square differences amongst the equivalent residues. The principal component 1 (PC1) graphic representation was displayed through Pymol Molecular Graphics System (https://pymol.org).

## 3 Results

### 3.1 Characterisation of N-terminal peptide substitutions that modulate coupling to Gα_s_ and β-arrestin-2 at GLP-1R and GCGR

The N-termini of GLP-1, glucagon and OXM play critical roles in the activation of their target receptors [31, 39]. However, the alanine (in GLP-1) or serine (in glucagon and OXM) at position 2 renders each of these endogenous ligands susceptible to DPP-4-mediated cleavage, and pharmacologically stabilised incretin analogues are often modified at this position. Here we focussed on the AIB2 substitution, found in semaglutide and some investigational oxyntomodulin analogues [40–42]. To systematically investigate how this change affects receptor activation, we obtained GLP-1-AIB2 and glucagon-AIB2 (GCG-AIB2) (see Table 1) and measured recruitment of β-arrestin-2 and mini-G_s_ to GLP-1R and GCGR in real time using nanoBiT complementation [25, 43]. Area-under-curve (AUC) quantification from kinetic response data indicated that efficacy for β-arrestin-2 recruitment to GLP-1R was modestly reduced with GLP-1-AIB2 compared to native GLP-1 (Table 2, Figure 1A, Supplementary Figure 1A). However, quantification of bias using the log(max/EC_50_) scale [26] indicated that this selective efficacy reduction does not qualify GLP-1-AIB2 as a biased agonist, as it was compensated by a corresponding small increase in potency (Figure 1B, 1C). The lack of bias is represented in Figure 1C by the 95% confidence intervals for GLP-1-AIB2 crossing zero. At GCGR, the impact of AIB2 was more striking, with large reductions in efficacy for both mini-G_s_ and β-arrestin-2 observed (Figure 1A); interestingly, this effect at GCGR could be partly reversed for both pathways by concurrent substitution of glutamine (Q) at position 3 to histidine (H), which our in-house preliminary evaluations had already flagged as a route to modulate GCGR signalling. GCG-AIB2 showed moderate but statistically significant degree of bias in favour of mini-G_s_ recruitment, with the H3 substitution driving the bias factor back towards zero (Figure 1B, 1C).

**Figure 1.**
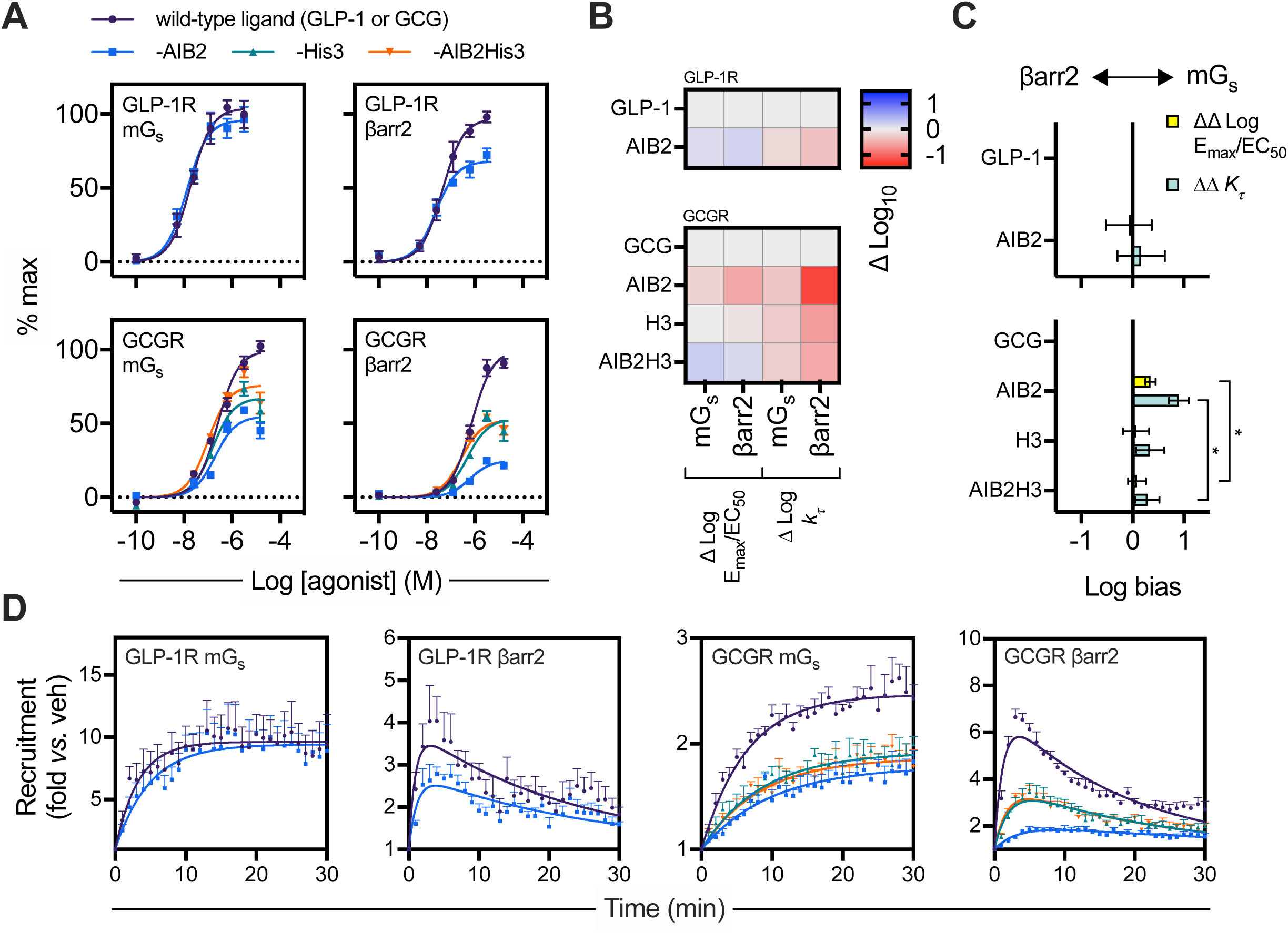
Evaluation of N-terminal substitutions to GLP-1, glucagon or OXM. (**A**) Concentration responses with 3-parameter fits showing mini-G_s_ (mG_s_) or β-arrestin-2 (βarr2) recruitment to GLP-1R-SmBiT or GCGR-SmBiT in HEK293T cells stimulated with GLP-1, GLP-1-AIB2, glucagon (GCG), GCG-AIB2, GCG-H3 or GCG-AIB2H3, *n*=5, with 3-parameter fits shown. (**B**) Heatmap representation of mean responses after quantification by log(max/EC_50_) or *k_τ_* method (see below) and normalisation to the reference ligand (GLP-1 or GCG, as appropriate). (**C**) Assessment of bias between mini-G_s_ and β-arrestin-2 recruitment from log(max/EC_50_) or *k*_*τ*_ methods, with statistical comparison by randomised block one-way ANOVA with Sidak’s test comparing GCG-AIB2 and GCG-AIB2H3. 95% confidence intervals are shown to allow identification of ligands with statistically significant bias *versus* the reference ligand. (**D**) Single maximal concentration kinetic responses for each ligand/receptor/pathway combination using data shown in (A), with one-phase association fits for mini-G_s_ and bi-exponential fits for β-arrestin-2. * p<0.05 by statistical test indicated. Data are represented as mean ± SEM for concentration response curves or 95% confidence intervals for bias plots; bias data are considered significant if the 95% confidence interval does not cross 0.

**Table 1.**
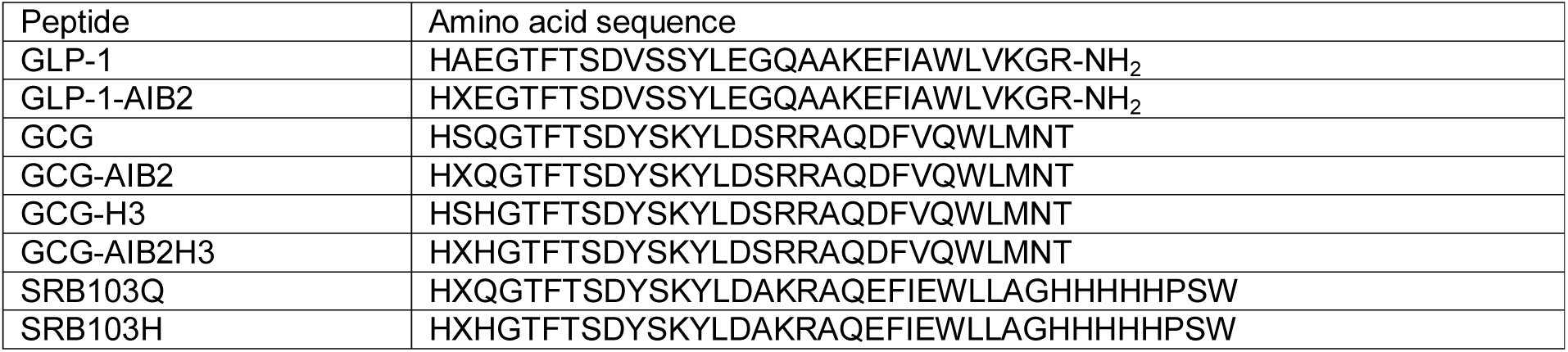
Amino acid sequences for peptides used in this study. Amino acid sequences are given in single letter code. GLP-1 is amidated at the C-terminus, as indicated. AIB is represented as “X”.

**Table 2.** Effect of AIB2 substitution in GLP-1, glucagon or OXM on mini-G_s_ and β-arrestin-2 recruitment responses. Mean ± SEM parameter estimates from 3-parameter fitting of AUC data from in Figure 1A, and association rate constants at maximal agonist stimulation (“*K*@[max]”). Statistical comparisons performed by paired t-test (GLP-1 and GLP-1-AIB2) or randomised block one-way ANOVA with Dunnett’s test (glucagon analogues). Note that, in general, if >1 ligand was a full agonist, E_max_ values were compared after normalisation to the globally fitted maximum response, whereas if only one ligand was a full agonist, statistical comparison was performed prior to normalisation, but numerical results are presented after normalisation to the full agonist response. See Supplementary Figure 1 for further analysis of β-arrestin-2 recruitment using a different system.* p<0.05 by indicated statistical test.

| GLP-1R |  |  |  |  |  |  |
| --- | --- | --- | --- | --- | --- | --- |
| | Mini-G<sub>s</sub> | | | β-arrestin-2 | | |
|  | pEC<sub>50</sub> (M) | E<sub>max</sub> (% max) | K@[max] (min<sup>-1</sup>) | pEC<sub>50</sub> (M) | E<sub>max</sub> (% max) | K@[max] (min<sup>-1</sup>) |
| GLP-1 | 7.7 ± 0.1 | 105 ± 2 | 0.30 ± 0.04 | 7.3 ± 0.2 | 100 ± 0 | 1.25 ± 0.31 |
| GLP-1-AIB2 | 7.9 ± 0.0 | 96 ± 2 | 0.21 ± 0.01 * | 7.7 ± 0.1 * | 68 ± 4 * | 0.82 ± 0.10 |
| GCGR |  |  |  |  |  |  |
| | Mini-G<sub>s</sub> | | | β-arrestin-2 | | |
|  | pEC<sub>50</sub> (M) | E<sub>max</sub> (% max) | K @[max] (min<sup>-1</sup>) | pEC<sub>50</sub> (M) | E<sub>max</sub> (% max) | K @[max] (min<sup>-1</sup>) |
| GCG | 6.7 ± 0.0 | 100 ± 0 | 0.17 ± 0.03 | 6.1 ± 0.1 | 100 ± 0 | 0.89 ± 0.18 |
| GCG-AIB2 | 6.7 ± 0.1 | 54 ± 4 * | 0.11 ± 0.01 | 6.2 ± 0.1 | 25 ± 2 * | 0.29 ± 0.03 * |
| GCG-H3 | 6.8 ± 0.1 | 66 ± 5 * | 0.13 ± 0.04 | 6.3 ± 0.1 | 54 ± 6 * | 0.51 ± 0.06 * |
| GCG-AIB2H3 | 7.0 ± 0.1 * | 74 ± 5 * | 0.13 ± 0.01 | 6.6 ± 0.1 * | 53 ± 2 * | 0.56 ± 0.06 * |

An alternative method for bias quantification has been proposed [27, 44] that is applicable to scenarios when kinetic response data is available. This model-free approach quantifies efficacy, termed k_*τ*_, from the initial response rate at a saturating agonist concentration. After logarithmic transformation of k_*τ*_, bias can be determined by first normalising to a reference ligand (giving Δlog k_*τ*_) and then comparing responses between pathways (giving ΔΔlog k_*τ*_). In our study, mini-G_s_ responses could be fitted as one-phase exponential association curves, whereas β-arrestin-2 showed a characteristic rapid increase and slower decline, presumed to reflect β-arrestin association followed by dissociation from the target receptor, and required a bi-exponential equation to define association and dissociation rate constants [45] (Figure 1D). GLP-1-AIB2 showed subtly slower kinetics at GLP-1R for both pathways than did GLP-1, which did not translate to a significant degree of bias using the ΔΔlog k*τ* method (Table 2, Figure 1C). At GCGR, mini-G_s_ and β-arrestin-2 association kinetics were also slower for GCG-AIB2 than for glucagon (Figure 1D, Table 2), with bias assessment from the kinetic data again suggesting a preference for mini-G_s_ coupling that was negated with the introduction of H3 (i.e. less bias with GCG-AIB2H3 than GCG-AIB2; Figure 1C).

Overall, these data indicate that introducing the AIB2 substitution into GLP-1 and glucagon leads to a noticeable reduction in efficacy for β-arrestin-2 recruitment, more than mini-G_s_ recruitment, with glucagon more affected than GLP-1. However, at GCGR, this effect could be mitigated by the presence of H3. The Q/H switch at position 3 thereby provides a means to modulate efficacy whilst retaining AIB2-induced resistance to DPP4.

### 3.2 GCGR molecular dynamics simulations

We performed molecular dynamics simulations of the active state GCGR in complex with glucagon, GCG-AIB2, or GCG-AIB2H3 to retrieve insights into the effects that peptide mutations have on the interactions, fingerprints and receptor flexibility. The substitution of serine at position 2 with the nonstandard residue AIB produced a substantial loss of interactions with the top of transmembrane helix 6 (TM6) and TM7 (E362^6.53^, F365^6.56^, and D385^7.42^ in Figure 2A, 2B). Fewer contacts were formed also with TM3 (I235^3.40^ and Y239^3.44^) and TM5 (W304^5.36^), compared to glucagon. The substitution of S2 with the hydrophobic AIB removed a persistent hydrogen bond with D385^7.42^ side chain (Table 3) and moved the barycenter of the interactions towards extracellular loop 2 (ECL2) (D299^ECL2^, S297^ECL2^ in Figure 2A, 2B) due to hydrogen bonds with H1 and T5 (Table 3). The partial release of TM6 from the restraining interactions with the peptide N-termini is corroborated by the high flexibility displayed in Figure 2C. GCGR in complex with glucagon and AIB2H3, on the other hand, was characterized by low plasticity of TM6, as indicated by monodisperse probability curves. Overall, glucagon and GCG-AIB2 stabilised divergent GCGR conformations of TM6, ECL2 and ECL3 (Figure 2D). Interestingly, in the closely related GLP-1R the ECL2 is essential for transducing peptide-receptor interactions into cAMP accumulation, while a possible correlation between peptides more prone to interact with ECL3 and β-arrestin-influenced signaling events such as ERK1/2 phosphorylation has been proposed [46].

**Figure 2.**
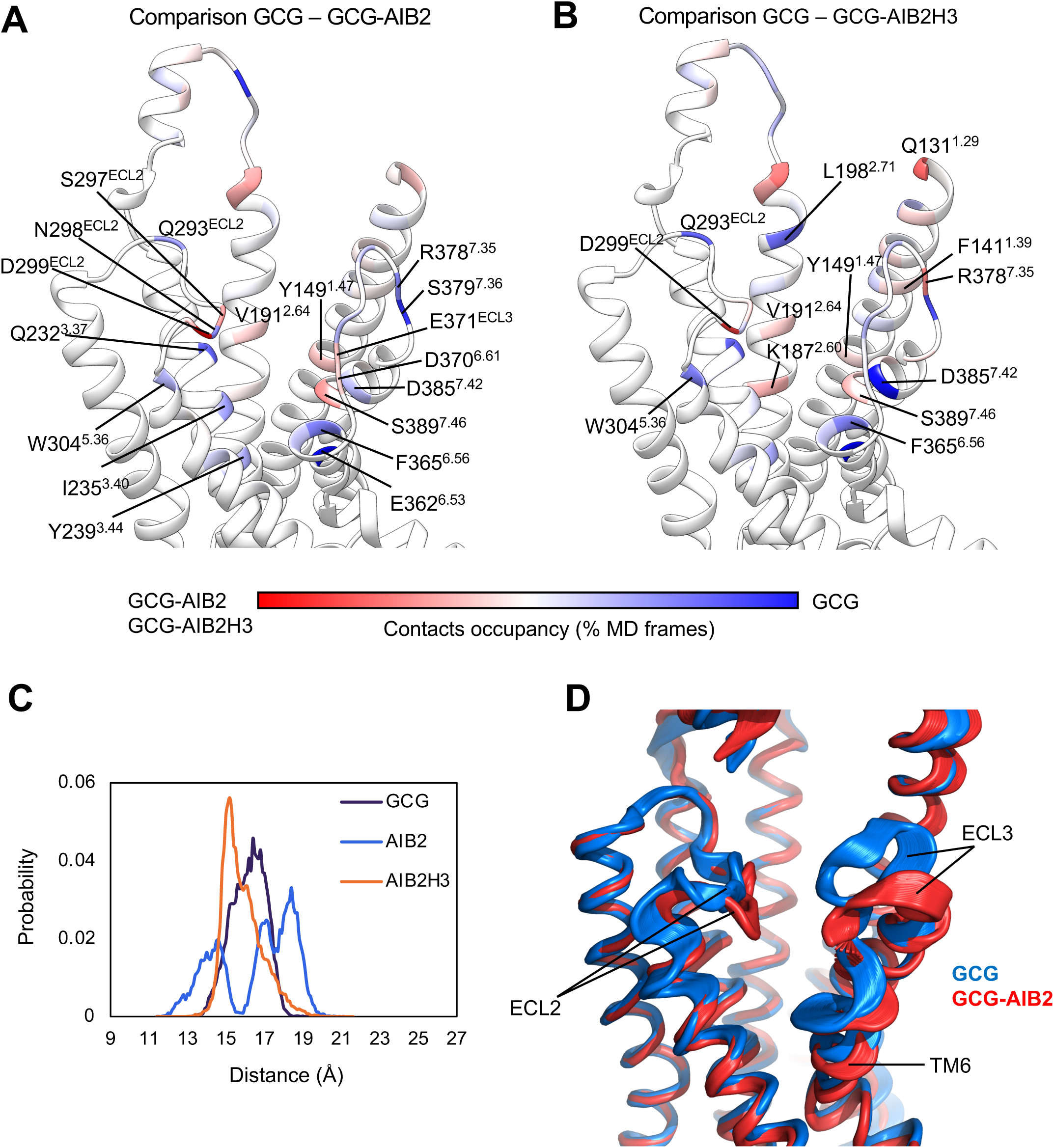
MD simulations of GCGR in complex with glucagon, GCG-AIB2, or GCG-AIB2H3. **A and B** show the difference in the contacts between GCGR and GCG-AIB2 (**A**) or GCG-AIB2H3 (**B**) plotted on the ribbon representation of GCGR; residues in red were more engaged by GCG-AIB2 (**A**) or GCG-AIB2H3 (**B**), while blue residues formed more contacts with GCG. **C)** Probability distribution of the distance between TM6 residue T369 and the origin of the cartesian coordinates (point 0,0,0). (**D**) Superposition of the PC1 analysis computed on the simulations of GCGR in complex with GCG (blue) or GCG-AIB2 (red).

**Table 3.** Molecular dynamics simulation results. Hydrogen bonds between GCGR and the first five amino acids in GCG, GCG-AIB2, and GCG-AIB2H3. Occupancy represents the number of frames with interaction divided by the total number of frames. ss indicate side chain – side chain hydrogen bonds, while sb indicate backbone – side chain hydrogen bonds.

| Peptide | Ligand residue | Receptor residue | Occupancy (%) | Type of hydrogen bond |
| --- | --- | --- | --- | --- |
| GCG | H1 | E362<sup>6.53</sup> | 37.4 | sb |
|  |  | D385<sup>7.42</sup> | 30.4 |  |
|  | S2 | D385<sup>7.42</sup> | 78.5 | ss |
|  |  |  | 18.4 | sb |
|  | Q3 | Y145<sup>1.43</sup> | 20.5 | ss |
|  |  | Y149<sup>1.47</sup> | 18.2 |  |
|  | T5 | N298<sup>ECL2</sup> | 10.6 | sb |
| GCG-AIB2 | H1 | D299<sup>ECL2</sup> | 30.2 | sb |
|  |  | K187<sup>2.60</sup> | 14.2 | ss |
|  |  | D385<sup>7.42</sup> | 11.7 |  |
|  | AIB2 | D385<sup>7.42</sup> | 43.2 | sb |
|  | Q3 | Y149<sup>1.47</sup> | 35.1 | ss |
|  |  | S389<sup>7.46</sup> | 13.6 |  |
|  | G4 | T296<sup>ECL2</sup> | 16.4 | sb |
|  | T5 | D299<sup>ECL2</sup> | 30.4 | ss |
| GCG-AIB2H3 | H1 | D299<sup>ECL2</sup> | 47.2 | sb |
|  |  | K187<sup>2.60</sup> | 2.46 | ss |
|  |  | N298<sup>ECL2</sup> | 14.1 |  |
|  | AIB2 | D385<sup>7.42</sup> | 21.0 | sb |
|  | H3 | Y149<sup>1.47</sup> | 11.9 | ss |
|  | T5 | D299<sup>ECL2</sup> | 13.8 |  |

Simulations suggested that the Q3H mutation introduced in GCG-AIB2H3 favors interactions between the peptide and TM2 residues K187^2.60^, V191^2.64^ and Q131^1.29^, located on the stalk region of the receptor. K187^2.60^, in particular, is part of the conserved hydrophilic region within class B receptor TMD, implied in binding, functionality and signal transmission [47]. It is plausible that the recovery in efficacy displayed by in GCG-AIB2H3 over AIB2 might be driven by stronger interactions with TM2. Moreover, the whole TMD closed up around GCG-AIB2H3 during simulations similarly to GCG (Figure 2C, 2D).

### 3.3 Pharmacologically stabilised GLP-1R/GCGR co-agonists to study the impact of efficacy variation

A pair of peptides termed “SRB103” (Table 1) was developed by an iterative process of sequence changes to the GLP-1R/GCGR co-agonist used in an earlier study [48]. As the previous peptide was derived from OXM, it contained the N-terminal sequence H-S-Q, which was modified to H-AIB-Q (SRB103Q) or H-AIB-H (SRB103H), along with additional conservative changes to enhance physicochemical properties such as stability and solubility. As expected, both SRB103Q and SRB103H were highly resistant to DPP-4-mediated degradation (Supplementary Figure 2A).

The mini-G_s_ and β-arrestin-2 recruitment profiles for each ligand were compared at both GLP-1R and GCGR (Figure 3A, Table 4, Supplementary Figure 2B). At GLP-1R, AUC analysis from the kinetic response data indicated a 40% reduction in β-arrestin-2 efficacy but a small increase in potency for the AIB2Q3 ligand compared to AIB2H3, with the mini-G_s_ response unaffected. At GCGR, both potency and efficacy were significantly reduced for both pathways with the AIB2Q3 ligand, although the magnitude of the efficacy reduction (∼20%) was small compared to the same sequence substitutions when applied to glucagon in Figure 1. Using the log(max/EC_50_) method there was no statistically significant bias between mini-G_s_ and β-arrestin-2 for SRB103Q *versus* SRB103H at either receptor (Figure 3B, 3C). However, bias estimates from the kinetic responses (ΔΔlog k_*τ*_ method) suggested a subtle preference for SRB103Q at GLP-1R towards mini-G_s_ recruitment (Figure 3C, 3D).

**Figure 3.**
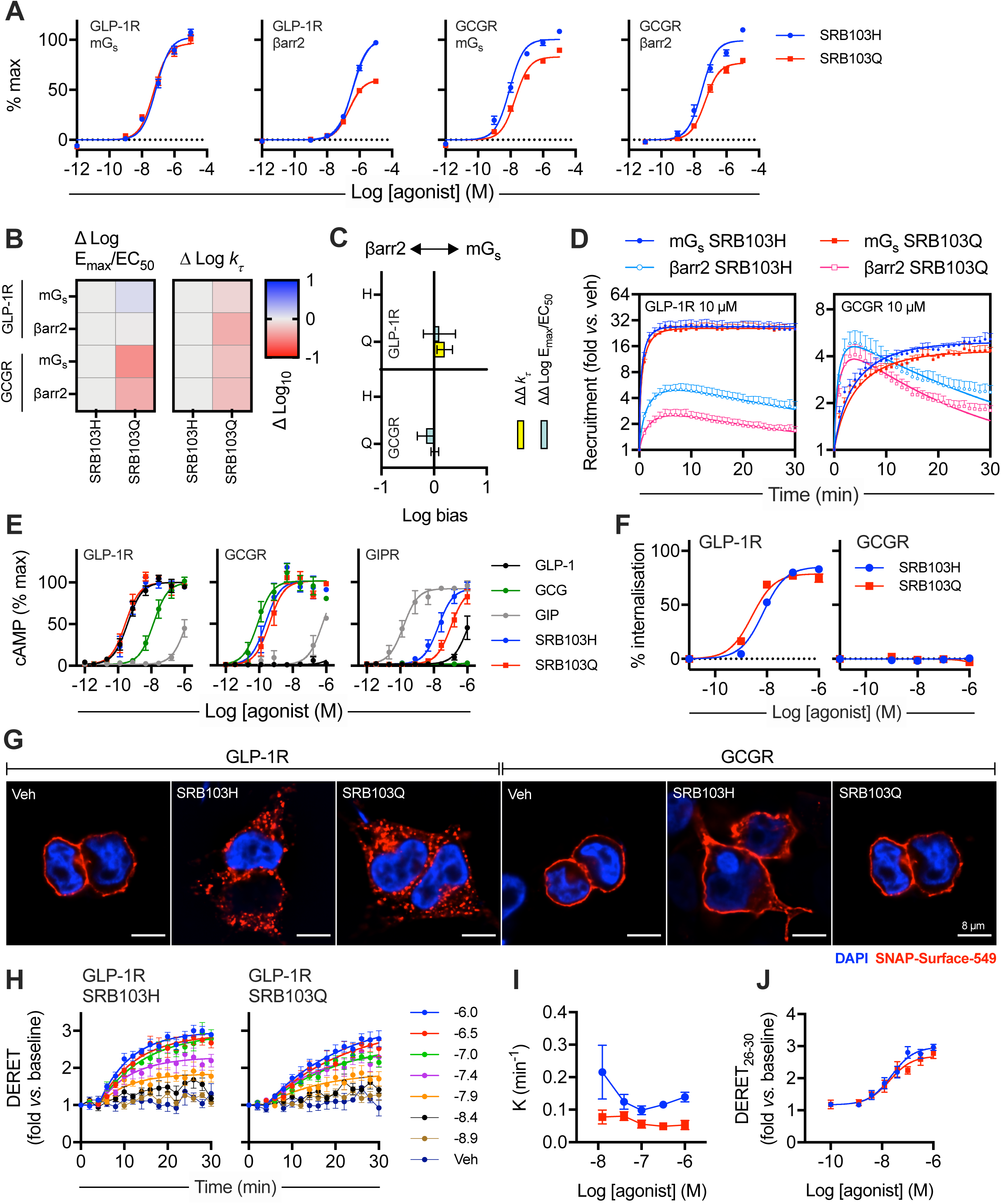
Development of a DPP-4-resistant GLP-1R/GCGR co-agonist with variable efficacy for intracellular effectors. (**A**) Concentration responses with 3-parameter fits for SRB103H- and SRB103Q-induced recruitment of mini-G_s_ or β-arrestin-2 to GLP-1R-SmBiT or GCGR-SmBit in HEK293T cells, *n*=6. (**B**) Heatmap representation of mean responses after quantification by log(max/EC_50_) or *k*_*τ*_method (see below) and normalisation to SRB103H as the reference ligand. (**C**) Assessment of bias between mini-G_s_ and β-arrestin-2 recruitment from log(max/EC_50_) or *k*_*τ*_ methods. 95% confidence intervals are shown to allow identification of ligands with statistically significant bias *versus* the reference ligand SRB103H. (**D**) Single maximal concentration kinetic responses for each ligand/receptor/pathway combination using data shown in (A), with one-phase association fits for mini-G_s_ and bi-exponential fits for β-arrestin-2. (**E**) cAMP responses in PathHunter CHO-K1 cells stably expressing GLP-1R, GCGR or GIPR, *n*=6, with 3-parameter fits shown. (**F**) SNAP-GLP-1R and SNAP-GCGR internalisation measured by high content analysis (HCA) in HEK293 cells, *n*=4, with 3-parameter fits shown. (**G**) Representative images from *n*=2 experiments showing endocytosis of SNAP-tagged receptors transiently expressed in HEK293 cells and treated with 100 nM agonist for 30 min. Scale bars = 8 µm. (**H**) SNAP-GLP-1R internalisation in HEK293-SNAP-GLP-1R cells, *n*=5, with one-phase association fits for ligand concentrations >10 nM shown (expressed as log[agonist] in M). (**I**) The concentration dependency of internalisation kinetics from (H) is shown. (**J**) Concentration responses are quantified from average response during the last 3 time-points from (H), with 3-parameter fits. Data are represented as mean ± SEM for concentration response curves or 95% confidence intervals for bias plots; bias data are considered significant if the 95% confidence interval does not cross 0.

**Table 4.** Pharmacological evaluation of SRB103H3 versus SRB103. Mean ± SEM parameter estimates from 3-parameter fitting of data from in Figures 3 and 4, and association rate constants for kinetic data where relevant. Statistical comparisons performed by paired t-test comparing SRB103Q *versus* SRB103H. If both ligands were full agonists, E_max_ values are shown after re-fitting of data normalised to the globally fitted maximum response. If only one ligand was a full agonist, statistical comparison was performed prior to normalisation, but numerical results are presented after normalisation to the full agonist response. * p<0.05 by indicated statistical test.

|  | SRB103H |  |  | SRB103Q |  |  |
| --- | --- | --- | --- | --- | --- | --- |
| | pEC50 (M) | $E_{\max}$ | $K@[max]$ ($\text{min}^{-1}$) | pEC50 (M) | $E_{\max}$ | $K@[max]$ ($\text{min}^{-1}$) |
| GLP-1R mini-$G_s$ (HEK293T) | $7.1 \pm 0.1$ | $103 \pm 4$ | $0.93 \pm 0.29$ | $7.3 \pm 0.1$ | $97 \pm 3$ | $0.70 \pm 0.20$ * |
| GLP-1R βarr2 (HEK293T) | $6.4 \pm 0.1$ | 100 | $0.37 \pm 0.06$ | $6.7 \pm 0.0$ * | $60 \pm 2$ * | $0.39 \pm 0.05$ |
| GCGR mini-$G_s$ (HEK293T) | $8.1 \pm 0.1$ | 100 | $0.14 \pm 0.01$ | $7.7 \pm 0.1$ * | $83 \pm 1$ * | $0.13 \pm 0.01$ |
| GCGR βarr2 (HEK293T) | $7.5 \pm 0.1$ | 100 | $0.81 \pm 0.07$ | $7.3 \pm 0.1$ * | $77 \pm 2$ * | $0.71 \pm 0.06$ * |
| GLP-1R cAMP (CHO-K1) | $9.5 \pm 0.1$ | $99 \pm 4$ | n.c. | $9.5 \pm 0.1$ | $103 \pm 2$ | n.c. |
| GCGR CAMP (CHO-K1) | $9.6 \pm 0.2$ | $107 \pm 5$ | n.c. | $9.4 \pm 0.2$ | $103 \pm 4$ | n.c. |
| GIPR CAMP (CHO-K1) | $7.6 \pm 0.2$ | $101 \pm 5$ | n.c. | $7.0 \pm 0.1$ * | $92 \pm 9$ | n.c. |
| DERET (HEK293-SNAP-GLP-1R) | $7.7 \pm 0.2$ | $3.0 \pm 0.1$ | $0.14 \pm 0.02$ | $7.7 \pm 0.2$ | $2.7 \pm 0.2$ * | $0.5 \pm 0.01$ * |
| HCA assay (HEK293-SNAP-GLP-1R) | $8.1 \pm 0.1$ | $86 \pm 2$ | | $8.6 \pm 0.0$ * | $79 \pm 3$ * | |
| cAMP 5 min (primary islet cells) | $9.0 \pm 0.4$ | $1.7 \pm 0.1$ | | $9.0 \pm 0.4$ | $1.6 \pm 0.1$ | |
| cAMP 10 min (primary hepatocytes) | $9.4 \pm 0.1$ | $170 \pm 7$ | | $8.9 \pm 0.1$ * | $175 \pm 12$ | |
| cAMP 16 h (primary hepatocytes) | $7.6 \pm 0.1$ | $206 \pm 7$ | | $7.5 \pm 0.0$ | $200 \pm 6$ | |

cAMP signalling responses were also assessed in CHO-K1 cell lines expressing GLP-1R, GCGR or glucose-dependent insulinotropic polypeptide receptor (GIPR) (Figure 3E, Table 4). Potencies for SRB103Q and SRB103H were, as expected, indistinguishable at GLP-1R, with a non-significant reduction for SRB103Q at GCGR. Both ligands showed at least 100-fold reduced potency for GIPR cAMP signalling compared to GIP itself, even in this highly amplified heterologous system, suggesting that GIPR is unlikely to contribute to their overall metabolic actions.

A close correlation has been observed previously between signalling efficacy and ligand-induced endocytosis of GLP-1R [19, 20]. GCGR, on the other hand, appears to internalise far more slowly [31, 49]. We investigated the effects of SRB103Q and SRB103H on internalisation of GLP-1R and GCGR SNAP-tagged at their N-termini in HEK293T cells by high content imaging [31]. Both ligands induced pronounced GLP-1R internalisation, with a minor reduction in efficacy with SRB103Q, but GCGR barely internalised with either ligand (Figure 3F); higher resolution images of SNAP-GLP-1R or SNAP-GCGR-expressing cells labelled prior to agonist treatment corroborated these findings (Figure 3G). Interestingly, when measured by diffusion-enhanced resonance energy transfer (DERET) [50], kinetics of GLP-1R internalisation were considerably slower for SRB103Q than for SRB103H throughout the concentration range (Figure 3H, 3I), although using AUC quantified from the end of the stimulation period, SRB103Q internalisation efficacy was only subtly reduced (Figure 3J), similar to the result in the high content imaging assay.

These data indicate that the AIB2Q3 iteration of RB103 shows reduced efficacy for recruitment of β-arrestin-2 at GLP-1R and, to a lesser degree, for mini-G_s_ and β-arrestin-2 at the GCGR.

### 3.4 Evaluating acute *versus* prolonged responses with SRB103H and SRB103Q

Reductions in β-arrestin-2 recruitment efficacy are associated with prolongation of cAMP signalling at GLP-1R [19, 39] and GCGR [31], thought to result from avoidance of target receptor desensitisation. In dispersed mouse islet cells, biochemically-measured acute cAMP responses to SRB103Q and SRB103H were indistinguishable (Figure 4A). Similarly, FRET imaging of intact mouse islets virally transduced to express the cAMP sensor epac2-camps [51] demonstrated that both agonists induce similar cAMP dynamics acutely (Figure 4B). However, when pre-treated for four hours with each ligand and then rechallenged after a washout period, a trend towards reduced responsiveness for SRB103H was observed. This difference was not significant when quantified from the whole re-stimulation period, but it was clearly observed that the epac2-camps average signal increase on SRB103H rechallenge was slower than for SRB103Q (*k*=0.28 *versus* 0.53 min^-1^ from pooled responses to SRB103H and SRB103Q, respectively), suggesting diminished responsiveness with the former ligand. GCGR responses were also evaluated in primary mouse hepatocytes; here, SRB103Q showed reduced potency acutely, but after overnight treatment, this difference had disappeared (Figure 4C). Overall, these studies indicate a general tendency for SRB103Q responses to be relatively enhanced with longer stimulations, which is compatible with reduced β-arrestin-mediated desensitisation of this ligand compared to SRB103H.

**Figure 4.**
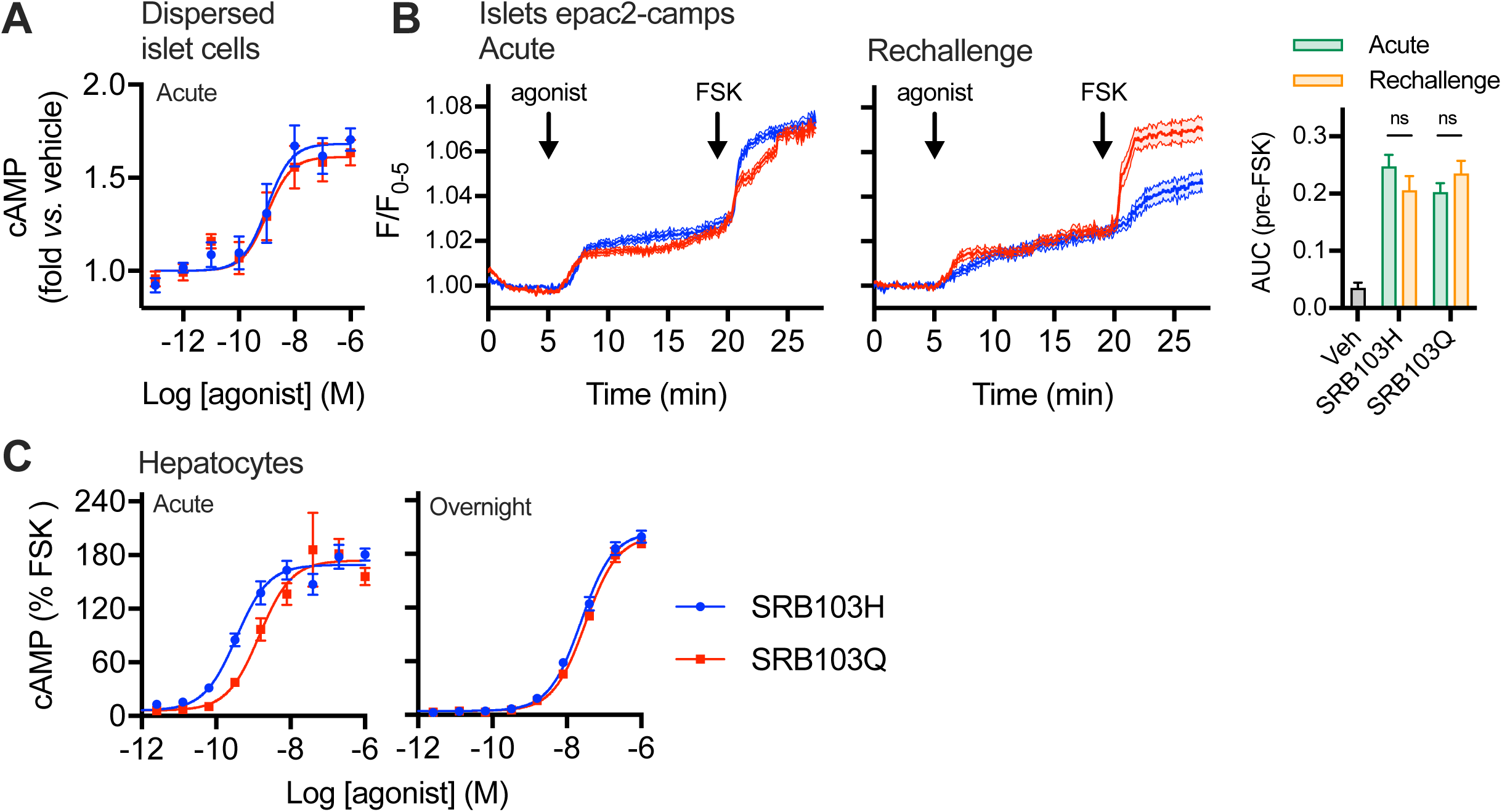
Acute *versus* prolonged responses *in vitro* with SRB103Q and SRB103H. (**A**) Acute cAMP signalling in primary dispersed mouse islets, 5 min stimulation with 500 µM IBMX, *n*=4, 3-parameter fits shown. (**B**) Whole islet cAMP responses to stimulation with 100 nM indicated agonist acutely or after 4-hour pre-treatment and washout, measured by FRET with virally transduced epac2-camps. Quantification from 25 – 42 mouse islets per treatment (5-9 mice from at least 2 independent islet preparations). AUCs during agonist exposure period (pre-forskolin [10 µM]) have been quantified and compared by two-way ANOVA with Sidak’s test. Representative images are shown. (**C**) Acute (10 min) and sustained (16 hour) cAMP accumulation in primary mouse hepatocytes, expressed relative to 10 µM forskolin response, *n*=4. Data are represented as mean ± SEM.

### 3.5 Anti-hyperglycaemic responses are prolonged after a single dose of SRB103Q *versus* SRB103H in mice

As GLP-1R agonists with reduced β-arrestin-2 recruitment efficacy and/or delayed endocytosis show progressive increases in anti-hyperglycaemic efficacy over longer exposure periods [19,21,52], we aimed to establish if this therapeutic principle could also be applied to GLP-1R/GCGR co-agonism. Indeed, blood glucose concentrations during an intraperitoneal glucose tolerance test (IPGTT) in lean mice tended to be lower after a single administration of SRB103Q compared to SRB103H, with this difference enhanced by longer agonist exposure time (Figure 5A). A similar pattern was seen at a range of agonist doses (Supplementary Figure 3A) and in diet-induced obese (DIO) mice (Figure 5B). Circulating concentrations of each ligand were the same 4 hours after a single dose, suggesting pharmacodynamic differences are unlikely to be due to altered pharmacokinetics (Supplementary Figure 3B).

**Figure 5.**
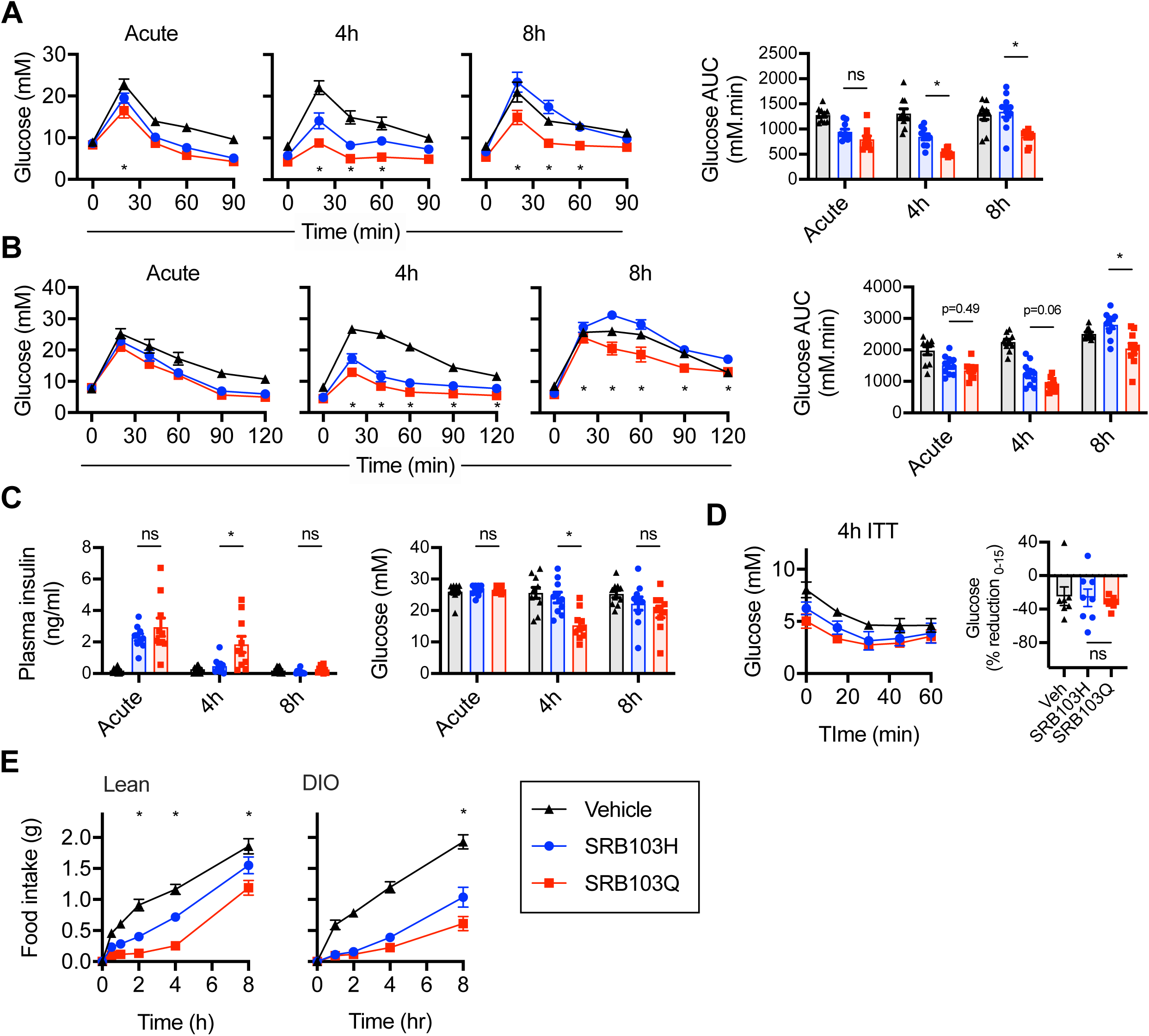
Immediate and delayed responses to SRB103Q and SRB103H in mice. (**A**) Blood glucose results during intraperitoneal glucose tolerance tests (IPGTTs) performed in lean male C57Bl/6 mice (*n*=10/group) with 2 g/kg glucose injected IP at the same time as, 4 hours after, or 8 hours after 10 nmol/kg agonist injection. Timepoint and AUC comparisons both by repeated measures two-way ANOVA with Tukey’s test; only SRB103Q *versus* SRB103H comparisons are shown. (**B**) As for (A) but in diet-induced obese male C57Bl/6 mice. (**C**) Plasma insulin and blood glucose results in lean male C57Bl/6 mice (*n*=10/group) 10 min after 2 g/kg IP glucose administration, concurrently with, 4 hours after or 8 hours after 10 nmol/kg agonist injection. AUC comparisons by repeated measures two-way ANOVA with Tukey’s test; only SRB103Q *versus* SRB103H comparisons are shown. (**D**) Blood glucose during insulin tolerance test (0.75 U/kg recombinant human insulin IP) performed 4 hours after administration of 10 nmol/kg agonist injection in lean male C57Bl/6 mice (*n*=8/group). Percentage reduction from 0 – 15 min is shown and compared by one-way ANOVA with Tukey’s test; only SRB103Q *versus* SRB103H comparison is shown. (**E**) Food intake in overnight-fasted lean male C57Bl/6 mice (*n*=8/group) treated with 10 nmol/kg indicated agonist. Timepoint comparisons both by repeated measures two-way ANOVA with Tukey’s test; only SRB103Q *versus* SRB103H comparisons are shown. * p<0.05 by indicated statistical test. Data are represented as mean ± SEM and with individual replicates where possible.

Both GLP-1R and GCGR agonism potentiate glucose-stimulated insulin secretion [12], but can also acutely enhance insulin-stimulated glucose disposal [11, 53]. Plasma insulin concentrations measured 10 min into a 4-hour-delayed IPGTT were higher with SRB103Q than SRB103H treatment, suggesting the former’s improved anti-hyperglycaemic efficacy is likely to derive at least partly from action on beta cells (Figure 5C), in keeping with the trend we observed in pancreatic islets towards improved responsiveness after prolonged stimulation with SRB103Q. In contrast, there was no evidence that insulin sensitivity was increased with either ligand, as assessed by insulin tolerance tests performed 4 hours after agonist administration (Figure 5D, Supplementary Figure 3C). Appetite suppression was also assessed in lean and diet-induced obese mice. Here, SRB103Q was more effective than SRB103H, particularly at later timepoints in the obese cohort (Figure 5E).

Overall, these results indicate that, despite showing lower acute efficacy for intracellular effector recruitment at both GLP-1R and GCGR, SRB103Q shows greater bioactivity in mice than SRB103H. For glycaemic effects, this difference tended to become more apparent with time, in keeping with the previously established principle that the metabolic advantages of biased GLP-1R agonists are temporally specific.

### 3.6 Improved anti-hyperglycaemic efficacy of SRB103Q is preserved with chronic administration

GLP-1R/GCGR co-agonists may hold advantages over GLP-1R mono-agonists for the treatment obesity and related metabolic diseases as their GCGR-mediated effects on energy expenditure can promote additional weight loss [33,54,55]. To determine if the apparent benefits of SRB103Q on glucose homeostasis revealed in single dose studies are also maintained after repeated dosing, SRB103H, SRB103Q and the GLP-1R mono-agonist liraglutide were administered at matched doses to DIO mice for 2 weeks. The dose was up-titrated over several days, analogous to typical practice in the clinic, as well as in preclinical studies of incretin receptor agonists [21, 56]. As expected, all agonists led to a significant amount of weight loss compared to vehicle (Figure 6A). However, the trajectory for weight lowering differed for both dual GLP-1R/GCGR agonists compared to liraglutide, with the latter being more effective earlier in the study before reaching a plateau after one week, as commonly observed with GLP-1R mono-agonists in rodents [57–59]. Importantly, the weight loss with both dual agonists was achieved despite liraglutide being more effective at suppressing energy intake throughout the study (Figure 6B), suggesting a contribution of increased energy expenditure [60]. Interestingly, SRB103Q was moderately more effective for weight loss than SRB103H despite similar energy intake, raising the possibility that reduced GCGR desensitisation could have contributed to improved longer term effects on energy expenditure. Both SRB103Q and SRB103H outperformed liraglutide in an IPGTT performed at the end of the study, with SRB103Q being the most effective at reducing the glucose excursion (Figure 6C).

**Figure 6.**
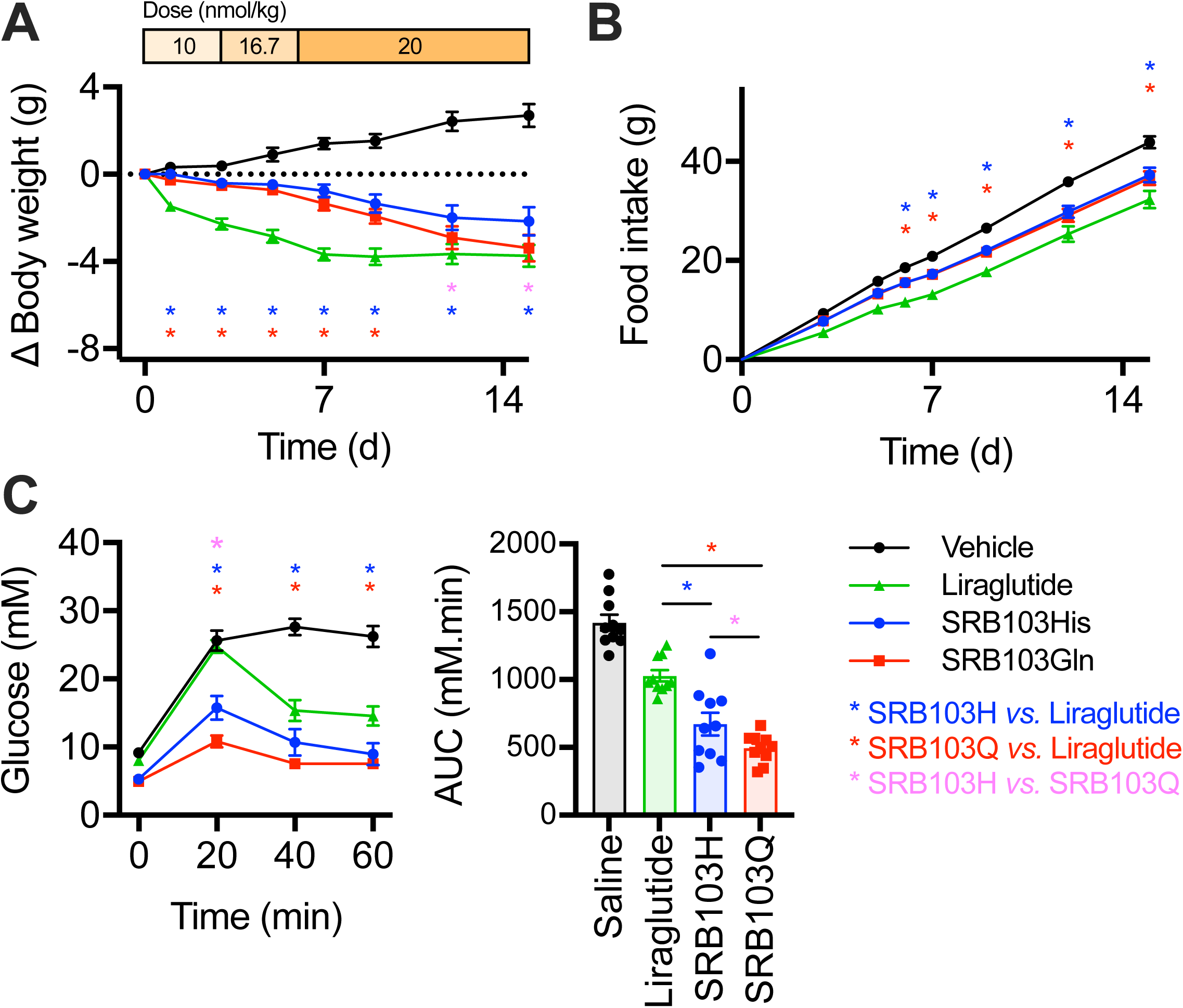
Repeated administration of SRB103Q and SRB103H. (**A**) Effect on body weight of daily administration by s.c. injection of SRB103Q, SRB103H, liraglutide or vehicle on body weight in male diet-induced obese C57Bl/6 mice, *n*=10/group with statistical comparisons between agonists by repeated measures two-way ANOVA with Holm-Sidak test. The injected daily dose is indicated above the graph. (**B**) As for (A) but cumulative food intake. (**C**) IPGTT (2 g/kg glucose) performed on day 15 of the study, 8 hours after agonist administration. Statistical comparisons were by repeated measures two-way ANOVA with Holm-Sidak test (time-points) or one-way ANOVA with Holm-Sidak test. * p<0.05 by indicated statistical test. Data are represented as mean ± SEM and with individual replicates where possible.

## 4 Discussion

In this study we have carefully evaluated the effects on GLP-1R and GCGR activity of the AIB2 substitution commonly used to confer DPP-4 resistance to therapeutic GLP-1R/GCGR peptide agonists. Depending on the peptide context, this substitution reduced efficacy for recruitment of key intracellular effectors at both target receptors. Interestingly, this effect was counteracted by substituting the neighbouring amino acid Q to H, providing a means to compare the impact of the resultant efficacy changes whilst retaining DPP-4 resistance. Whilst the efficacy-reducing effect of AIB2 was most prominently observed with glucagon itself at GCGR, in the context of the SRB103 peptides this effect was in fact greater at GLP-1R, specifically for β-arrestin-2 recruitment, although GCGR responses were also modestly reduced. The potential importance of this pharmacological finding was hinted at by studies in primary islets and hepatocytes – tissues in which these responses are chiefly driven by, respectively, GLP-1R and GCGR – where we observed that the lower efficacy SRB103Q ligand showed trends towards relatively enhanced signalling responses at both GLP-1R and GCGR over time. These observations support our *in vivo* findings that the improved anti-hyperglycaemic performance of SRB103Q becomes more apparent at later time-points after dosing, as was previously seen with GLP-1R mono-agonists with analogous signalling parameters [19].

This study was originally designed to assess the potential for biased agonism to improve therapeutic targeting of GLP-1R and GCGR. However, the magnitude of bias between SRB103Q and SRB103H, as assessed by two validated models, was relatively small. Interestingly, whilst biased agonism has attracted much attention over recent years, it has also been suggested that low intrinsic signalling efficacy, rather than biased agonism *per se*, is a viable alternative explanation for the improved performance of certain µ opioid receptor agonists [61], a GPCR target usually considered highly tractable to biased agonism [62]. This possibility is reinforced by a lack of consistency between formal bias estimates obtained from different analytical approaches, which can lead to different conclusions from the same data [63]. With regard to the lower efficacy SRB103Q agonist in our study, signal amplification downstream of Gα_s_ activation means full cAMP/PKA responses are still possible, so, in combination with reduced efficacy for β-arrestin recruitment, this could lead to reductions in desensitisation over time and allow longer-lasting signalling responses. Thus, beneficial responses from partial agonism may be achieved irrespective of whether formally quantified bias is present or not. Further evaluations to establish whether partial agonism or bias is the most important factor will be required to settle this issue.

The AIB2 substitution at position 2 is one of a number of sequence modifications that has been trialled to obtain DPP-4 resistance for incretin receptor analogues. Whilst exendin-4, the prototypical DPP-4 resistant GLP-1R mono-agonist, contains a glycine at position 2, GLP-1-G2 was recognised in early studies to show an unacceptable loss of signalling potency [64]; more recently it was demonstrated that this is also associated with reduced efficacy for recruitment of both mini-G_s_ and β-arrestin-2 to the GLP-1R [39]. AIB2 is better tolerated by GLP-1 than G2, whilst retaining identical protection against DPP-4 mediated degradation [64], and has been incorporated into the current leading GLP-1R mono-agonist semaglutide [65]. Our new data indicates that, in the context of native GLP-1, AIB2 leads to a significant reduction in efficacy for recruitment of β-arrestin-2 whilst barely affecting recruitment of mini-G_s_. This effect is likely to be peptide-specific, as we did not observe similar reductions in β-arrestin recruitment by AIB2-containing semaglutide in a previous study [19]. Interestingly, in the present work, AIB2 led to marked attenuation of engagement of GCGR with intracellular effectors by glucagon analogues, an effect which was previously hinted by the lower cAMP signalling potency with a glucagon analogue bearing AIB at both position 2 and 16 [66]. In the latter study, GCGR signalling was partly restored by conjugation to a fatty acid moiety, a well-established strategy used primarily to extend peptide pharmacokinetics by promoting reversible binding to albumin but, in this case, also found to enhance receptor activation. In our study we observed that switching Q to H at position 3 of glucagon was an alternative means to reverse the deleterious effect of AIB2 on GCGR signalling. It is not clear however if these strategies are equivalent, as in the study of Ward *et al*. [66] the signalling deficit seen with AIB2 was a reduction in cAMP potency, whereas in our study efficacies for mini-G_s_ and β-arrestin-2 was reduced but potencies were unaffected. A recent evaluation of GLP-1R/GCGR co-agonists [22] showed that the GLP-1R/GCGR/GIPR “tri-agonist” (GLP-1R/GCGR/GIPR) peptide originally described by Finan *et al*. [67], which includes the N-terminal sequence H-AIB-Q, does indeed show reductions in β arrestin recruitment efficacy to GLP-1R (modest) and GCGR (substantial) compared to the endogenous agonist, without major loss in potency, broadly matching our observations with native ligand analogues and SRB103 peptides. Measured signalling potency, especially in the context of significantly amplified responses e.g. cAMP, is driven to varying extents by both affinity and efficacy, with our results highlighting how the standard approach to evaluate incretin receptor agonists *in vitro* using cAMP in heterologous systems, which tends to render all compounds full agonists, may be insufficient to adequately decipher ligand pharmacology [68]. Importantly, our study also provides structural insights into the importance of position 2 of glucagon peptide analogues, with molecular dynamics simulations indicating that the reduced β-arrestin-2 recruitment associated with the AIB2 substitution is related to reduced engagement of ECL3.

The most striking results in our study were observed from *in vivo* comparisons of SRB103Q and SRB103H. Here, the lower efficacy SRB103Q (at both GLP-1R and GCGR) peptide outperformed SRB103H for ability to lower blood glucose levels 4 and 8 hours after a single injection, despite identical pharmacokinetics. These findings are reminiscent of observations with exendin-phe1, a GLP-1R mono-agonist with marked reductions in β-arrestin recruitment efficacy, which displayed better anti-hyperglycaemic effects and increased insulin secretion compared to exendin-4 in mice, with these differences being most obvious at later time-points [19]. However, one of the challenges with our study is identifying whether the observed effects result from enhanced action primarily at GLP-1R or at GCGR, as SRB103Q displayed reduced β-arrestin-2 recruitment efficacy at both receptors, meaning that longer lasting signalling through avoidance of target desensitisation could apply in both cases. Overall, we favour a primarily GLP-1R-mediated mechanism because 1) the selective reduction in β-arrestin-2 recruitment with SRB103Q was larger at GLP-1R than at GCGR, and 2) the effect was associated with increases in insulin release and supported by a trend towards reduced islet desensitisation *in vitro*. Whilst glucagon can augment glucose-stimulated insulin secretion, this effect is mediated mainly by cross-reactivity at the GLP-1R [12]. We consider it unlikely that the lower blood glucose levels with SRB103Q result from decreased hepatic glucose output via the subtly reduced efficacy of this peptide at GCGR, as it retains full cAMP activity in mouse hepatocytes, especially with prolonged stimulations, and the observed glycaemic effects relate mainly to the ability to restrain the hyperglycaemic effect of exogenously administered glucose. As antagonists for GLP-1R and GCGR are generally unable to cleanly and completely inhibit the action of high affinity exogenously administered agonists at pharmacological doses, studies in GLP-1R and GCGR knockout mice will be needed to distinguish the relative contributions of each receptor. The well-known phenotype of GCGR knockout mice, which are highly resistant to hyperglycaemia and show other metabolic abnormalities [69], may introduce additional challenges however.

SRB103Q and SRB103H were compared in a chronic administration study, with liraglutide also included for reference as an exemplar GLP-1R mono-agonist. The important observation here was that the enhanced anti-hyperglycaemic benefits of SRB103Q were retained after 2 weeks of repeated administration, suggesting that the apparent benefits of its intracellular signalling profile on glucose homeostasis do not diminish with time. This represents, to our knowledge, the first demonstration of the possibility of achieving more effective metabolic control through partial agonism in the context of a GLP-1R/GCGR co-agonist. Notably, somewhat greater weight loss without a corresponding reduction in food intake for SRB103Q was observed, which could conceivably result from increases in sustained GCGR activation, as might be predicted from the reduced β-arrestin-2 recruitment efficacy of this peptide compared to SRB103H. The glycaemic effects of both molecules compared favourably with liraglutide at the same dose, although differences in pharmacokinetics (longer with liraglutide) and the amount of bioactive free peptide (lower with liraglutide due to albumin binding) complicate interpretation. Nevertheless, the observation that both SRB103 peptides achieve similar weight loss to liraglutide despite a less potent anorectic effect adds to the evidence that GLP-1R/GCGR co-agonism may be an effective means to treat obesity, potentially with reduced anorexia-associated nausea (although this was not tested directly in our study).

The recent study of Darbalaei *et al*. provides a comprehensive description of the pharmacology of other published GLP-1R/GCGR co-agonists [22], including two ligands for which clinical data are available – cotadutide (MEDI0382) [8] and SAR425899 [70]. Neither of these clinical candidate molecules include AIB2 at position 2, but both showed reduced recruitment of β-arrestin-2 to GLP-1R, albeit the reduction was not as great as for SRB103Q. Both also showed significantly reduced recruitment of β-arrestin-2 to GCGR compared to glucagon, a difference that was larger than the GCGR efficacy reduction seen with SRB103Q compared to SRB103H. Thus, cotadutide and SAR425899 may well be additional examples of incretin receptor ligands retrospectively identified as showing biased agonist properties, as was recently found for the dual GLP-1R/GIPR agonist tirzepatide [71, 72]. On the other hand, the recorded cAMP potencies for cotadutide and SAR425899 in the study of Darbalaei *et al*. relative to the endogenous comparator ligands are orders of magnitude less than reported previously [70, 73], raising the possibility that the cellular systems used to evaluate the pharmacology of these ligands could have affected the results.

In conclusion, our study should be seen as an evaluation of the potential for reduced efficacy to be incorporated into the assessment process for candidate dual GLP-1R/GCGR agonists. Further molecular optimisations, e.g. acylation for extended pharmacokinetics, will be required to generate viable molecules for eventual clinical use. Molecular dynamics simulations have indicated relevant differences in engagement with ECL2 and ECL3 that can be used to guide these optimisations. Detailed mechanistic work is also needed to establish the relative contributions of G protein and β arrestin-mediated effects at both GLP-1R and GCGR, and will help clarify how investigational incretin receptor agonists are prioritised during drug development for T2D and obesity.

## Funding acknowledgements

The Section of Endocrinology and Investigative Medicine is funded by grants from the MRC, BBSRC, NIHR, and is supported by the NIHR Biomedical Research Centre Funding Scheme. The views expressed are those of the authors and not necessarily those of the funders. The views expressed are those of the authors and not necessarily those of the funder. D.J.H. was supported by MRC (MR/N00275X/1 and MR/S025618/1) and Diabetes UK (17/0005681) Project Grants. This project has received funding from the European Research Council (ERC) under the European Union’s Horizon 2020 research and innovation programme (Starting Grant 715884 to D.J.H.). A.T. acknowledges funding from Diabetes UK. B.J. acknowledges support from the Academy of Medical Sciences, Society for Endocrinology, The British Society for Neuroendocrinology, the European Federation for the Study of Diabetes, and an EPSRC capital award. B.J. and A.T. also received funding from the MRC (MR/R010676/1) and the European Federation for the Study of Diabetes. E.R.M. acknowledges support from Royal College of Surgeons of England and MRC clinical research training fellowships.

## Abbreviations

AIB: 2-aminoisobutyric acid
βarr: β-arrestin
cAMP: Cyclic adenosine monophosphate
DERET: Diffusion-enhanced resonance energy transfer
DPP-4: Dipeptidyl dipeptidase-4
ECL: Extracellular loop
GCG(R): Glucagon (receptor)
GIP(R): Glucose-dependent insulinotropic polypeptide (receptor) GLP-1(R) Glucagon-like peptide-1 (receptor)
HCA: High content analysis
IPGTT: Intraperitoneal glucose tolerance test
OXM: Oxyntomodulin
PKA: Protein kinase A
T2D: Type 2 diabetes
TM: Transmembrane (helix)

## Acknowledgements

We are grateful to Zainab Malik for technical assistance.

