## Supplementary Information for "Partial agonism improves the anti-hyperglycaemic efficacy of an oxyntomodulin-derived GLP-1R/GCGR co-agonist"

**Contents:**

- **Supplementary Figure 1**
- **Supplementary Figure 2**
- **Supplementary Figure 3**

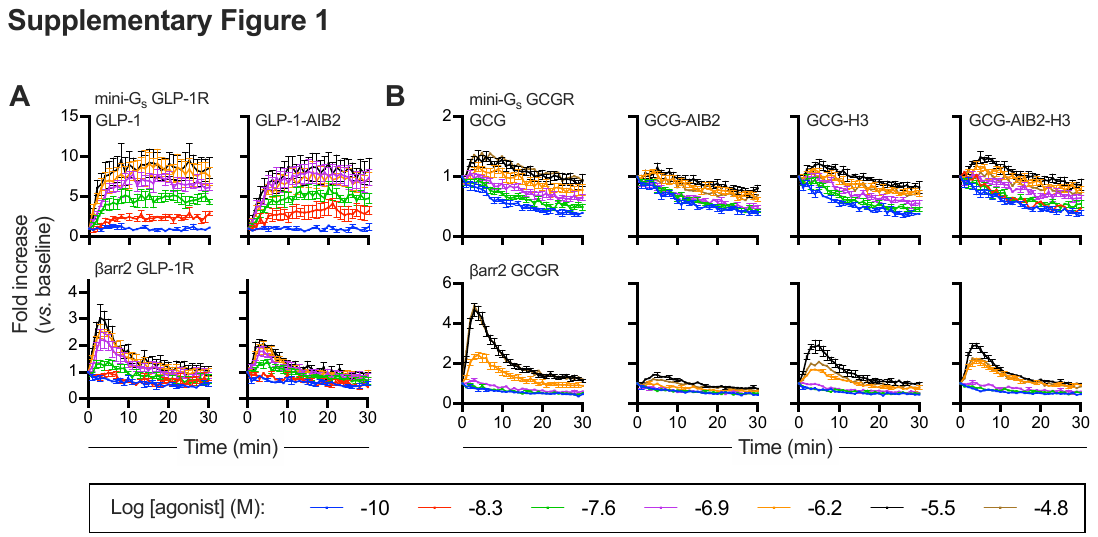

**Supplementary Figure 1. Nanobit kinetic traces with GLP-1 and glucagon analogues.** (**A**) Recruitment of mini-G_s_ or β-arrestin-2 to GLP-1R-SmBiT in HEK293T cells, responses to GLP-1 and GLP-1-AIB2, *n*=5. (**B**) As for (A) but for GCGR-SmBiT and GCG, GCG-AIB2, GCG-H3 and GCG-AIB2H3. Mean ± SEM responses are shown.

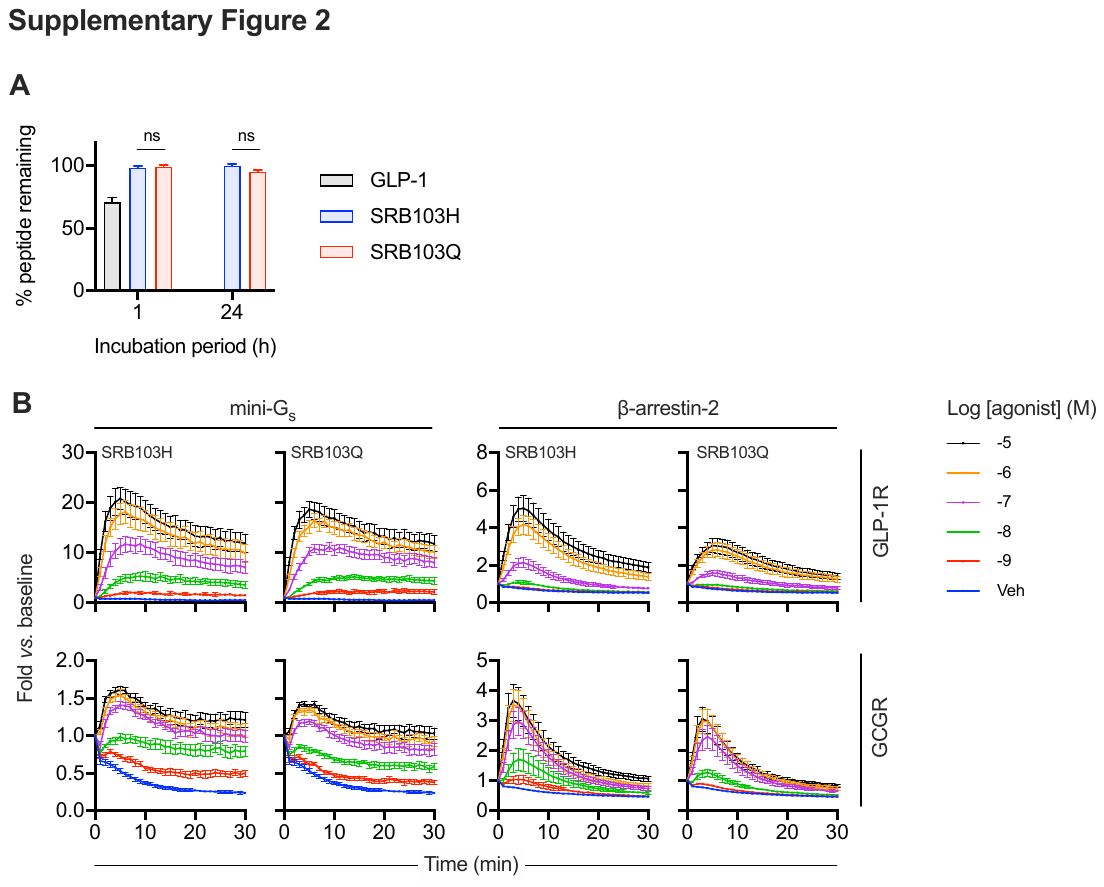

**Supplementary Figure 2. Further data for SRB103 analogues.** (**A**) Assessment of DPP-4 sensitivity for each peptide by HPLC measurement of intact peptide remaining after incubation with recombinant DPP-4 at 37°C for 1 or 24 hours, *n*=3, compared by repeated measures two-way ANOVA with Tukey’s test; only SRB103Q *versus* SRB103H comparisons are shown. (**B**) Recruitment of mini-G_s_ or β-arrestin-2 to GLP-1R-SmBiT or GCGR-SmBiT in HEK293T cells, responses to SRB103 peptides, *n*=6. Mean ± SEM responses are shown.

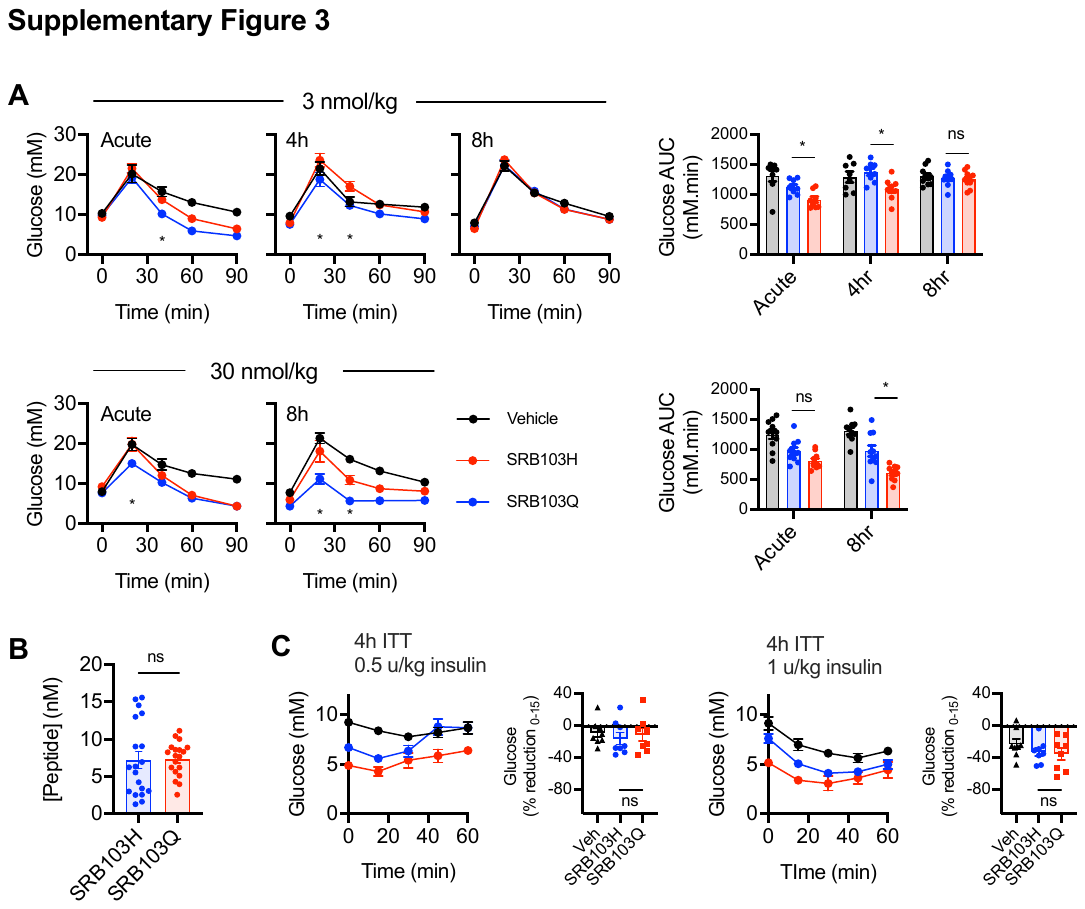

**Supplementary Figure 3. SRB103 analogue responses in mice.** (**A**) Blood glucose results during intraperitoneal glucose tolerance tests (IPGTTs) performed in lean male C57Bl/6 mice (*n*=8-10/group) with 2 g/kg glucose injected IP at the same time as, 4 hours after, or 8 hours after 3 or 30 nmol/kg agonist injection. Timepoint and AUC comparisons both by repeated measures two-way ANOVA with Tukey’s test; only SRB103Q *versus* SRB103H comparisons are shown. (**B**) Plasma concentration of SRB103H and SRB103Q 4 hours after IP injection of 0.5 mg/kg agonist, as determined by radioimmunoassay, compared by two-tailed unpaired t-test. (**C**) Blood glucose during insulin tolerance tests (0.5 or 1 U/kg actrapid insulin IP) performed 4 hours after administration of 10 nmol/kg agonist injection in lean male C57Bl/6 mice (*n*=8/group). Percentage reduction from 0 – 15 min is shown and compared by one-way ANOVA with Tukey’s test; only SRB103Q *versus* SRB103H comparisons are shown. * p<0.05 by indicated statistical test. Data are represented as mean ± SEM and with individual replicates where possible.
